# A geometric characterization of population coding in the prefrontal cortex and hippocampus during a paired-associate learning task

**DOI:** 10.1101/578849

**Authors:** Yue Liu, Scott L. Brincat, Earl K. Miller, Michael E. Hasselmo

**Affiliations:** Department of Physics, Boston University, Boston, MA 02215; Center for Systems Neuroscience, Boston University, Boston, MA 02215; Department of Psychological and Brain Sciences, Boston University, Boston, MA 02215; The Picower Institute for Learning and Memory, Massachusetts Institute of Technology, Cambridge, MA 02139; Department of Brain and Cognitive Sciences, Massachusetts Institute of Technology, Cambridge, MA 02139

## Abstract

Large-scale neuronal recording techniques have enabled discoveries of population-level mechanisms for neural computation. However it is not clear how these mechanisms form by trial and error learning. In this paper we present an initial effort to characterize the population activity in monkey prefrontal cortex (PFC) and hippocampus (HPC) during the learning phase of a paired-associate task. To analyze the population data, we introduce the normalized distance, a dimensionless metric that describes the encoding of cognitive variables from the geometrical relationship among neural trajectories in state space. It is found that PFC exhibits a more sustained encoding of task-relevant variables whereas HPC only transiently encodes the identity of the stimuli. We also found partial evidence on the learning-dependent changes for some of the task variables. This study shows the feasibility of using normalized distance as a metric to characterize and compare population level encoding of task variables, and suggests further directions to explore the learning-dependent changes in the population activity.

## 1 Introduction

With the development of experimental techniques for recording the activity of a large number of neurons, researchers have started exploring the possibility that the collective dynamics of interacting populations of neurons forms basic units for some neural computations (Sussillo (2014)). The collective dynamics are usually described by neural trajectories in state space which represent firing rates evolving through time. By looking at the collective neural dynamics through the lens of dynamical systems, several studies have identified cognitive functions with familiar concepts in dynamical systems. For example, in Mante et al. (2013), it was shown that the monkey prefrontal network performed a context-dependent decision making task by forming a pair of line attractors for the two contexts (Mante et al. (2013)). In Remington et al. (2018), the authors showed that in an interval production task, monkey frontal cortex circuits encoded the information about the time interval to be reproduced in the initial condition of the neural population dynamics, and the neural dynamics for different reproduced time intervals were represented by parallel neural trajectories with different speeds (Remington et al. (2018)).

Despite great progress in revealing collective functional features in neural computation, it remains unclear how these collective features are formed during training. Although it has been shown that similar features are present in recurrent neural networks (RNNs) trained on the same task by backpropagation, the learning dynamics of these RNNs are likely to differ from real neural circuits. To elucidate the solutions to this problem, one should look at how population dynamics change during the learning phase of a task. There exists studies that look at population level changes during motor learning (Sadtler et al. (2014); Golub et al. (2018); Vyas et al. (2018)), but similar work for cognitive learning has been scarce (although see Durstewitz et al. (2010)). In this paper we present an initial effort to characterize population level dynamics during the learning phase of a cognitive task.

The task we analyzed is a paired associate task where a monkey learned associations between a pair of randomly chosen visual stimuli (cues) and a third visual stimulus (associate). We are interested in the type of changes in the population activity that correlate with learning in this task. The ways the network learns to solve this task could be broadly divided into two categories. It could be that the network develops more similar representations for the cues that belong to the same associate, as previously suggested by single-cell analysis (Brincat and Miller (2016)) as well as results showing categorical representation in prefrontal cortex (e.g. Freedman et al. (2001); Roy et al. (2010)). Another possibility is that the network response to cues does not change with learning, but the cue leaves a trace of lasting changes in the network state. The network then learns to combine its state at the presentation of the associate with the identity of the associate to generate the appropriate decision. In this scenario, learning is not necessarily reflected in the change of neural activity but could be some “silent” mechanisms such as synaptic mechanisms. As will be shown, the analysis in the paper suggests the second possibility.

For analysis of the neural recording data, we introduced a dimensionless metric which we called normalized distance (ND) that describes the geometric relationships among neural trajectories for different experimental conditions. Unlike decoding methods, ND characterizes the encoding of all task variables in the population based directly on the geometry of the neural code. The dimensionless property of ND also enables comparisons of information in population codes between different learning stages as well as different brain areas. Using this metric, we then compared population level dynamics between prefrontal cortex (PFC) and hippocampus (HPC) as well as across learning stages in terms of the information content in the population codes. Our results reveal a series of differences in the dynamics of the information content between PFC and HPC. Differences in population coding across learning stages are also present, albeit in one of the two animals. These results demonstrate that normalized distance is a robust way of measuring information content in population codes with high temporal resolution in the face of noisy neural data.

## 2 Methods

### 2.1 Task and recording

We analyzed neural recording data from a previous study on a pair-associate learning task (Brincat and Miller (2015)). In that study, two macaque monkeys were trained to perform an object paired-associate learning task that required them to learn arbitrary associations between pairs of visual images of objects. On each day, six novel images were chosen. Four of them were randomly assigned as the cue objects and the other two were assigned as the associate objects. Two random cue objects were then paired with a random associate object, forming a 4:2 mapping from cues to associates (Figure 1a).

**Figure 1:**
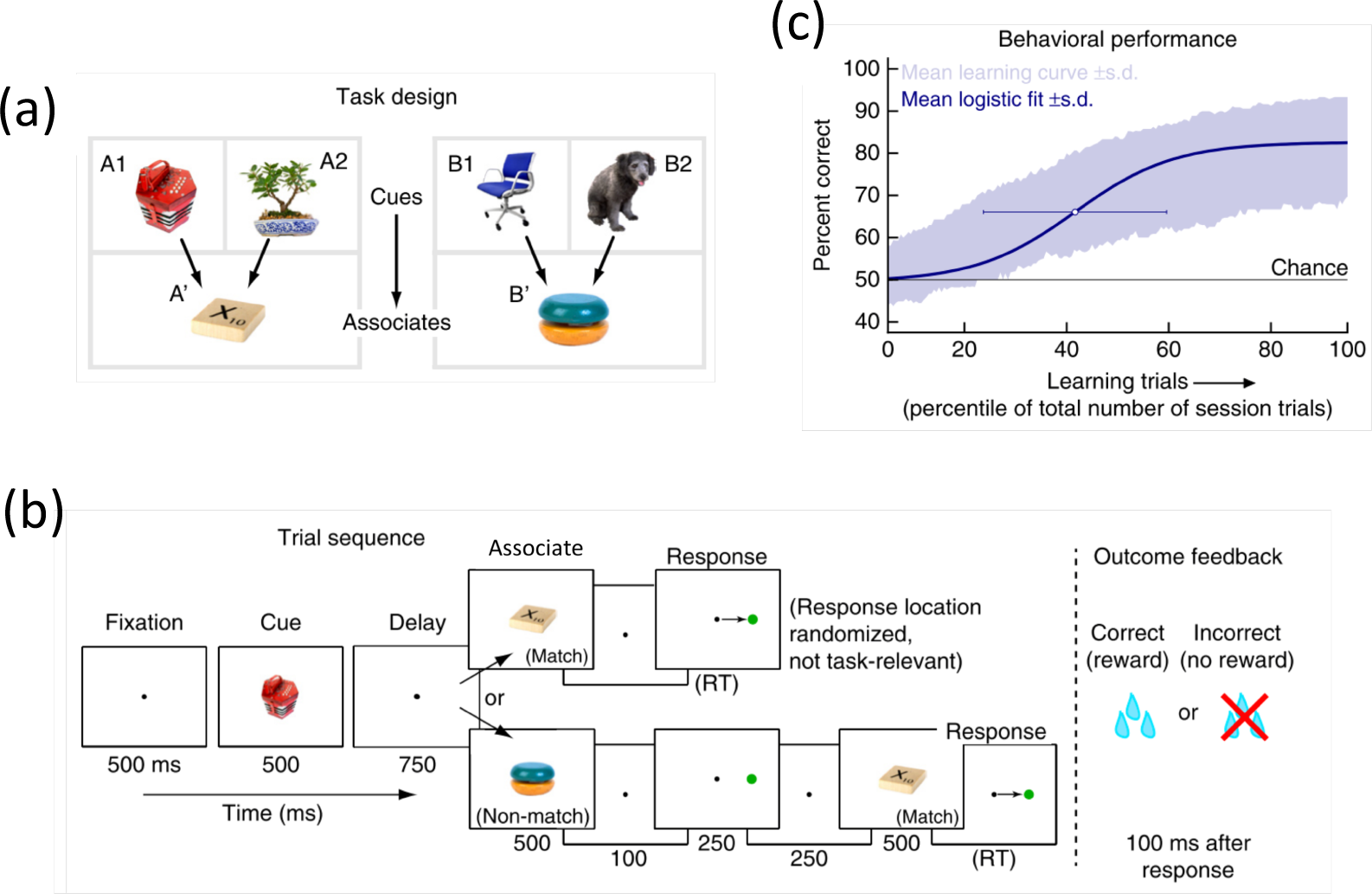
Overview of the task. On each session, six random stimuli are arranged into two-to-one mappings (**a**) and the monkey is required to learn the mappings via trial and error. On each trial, the monkey receives a cue and a choice stimuli separated by a 750ms delay, after which it has to make a decision of whether the two stimuli match by a saccade response for match. For nonmatch trials, the match stimulus is shown after 250 ms after which the monkeys are required to make the same saccade as in match trials (**b**). During each session the monkey’s performance gradually increases (**c**). Adapted from Brincat et al., 2015.

The structure of a trial is illustrated in Figure 1b. During each trial, after a 500 ms fixation period, the monkeys were first presented with one of the cue stimuli for 500 ms. Then, after a delay of 750 ms, the monkeys were presented with one of the associate objects for another 500 ms. The monkeys should indicate whether the previous two stimuli are associated with each other by making a saccade to the indicated position on the screen if they match. If the two previous stimuli do not match, the monkeys were required to hold the fixation. After a 250 ms delay, the match stimulus will appear on the screen for 500 ms after which the monkeys were required to make the saccade to the same position as in the match trial. The monkeys were rewarded with water if the response was correct (Figure 1b).

During each recording session, the monkey must learn novel associations from trial and error. The monkey was able to learn the associations above chance in all recording sessions (Figure 1c). Microelectrodes were lowered into the lateral prefrontal cortex and hippocampus and recorded spike and LFP signals while the monkeys were performing the paired-associate learning task. Across all sessions, a total of 496 neurons in prefrontal cortex 270 neurons in hippocampus were recorded. For each session, 6-134 neurons were simultaneously recorded in PFC and 4-134 in HPC.

### 2.2 Normalized distance

In this section we introduce the metric we used to characterize the information content in a neural population. This metric is computed from distance between neural trajectories in state space, but normalized properly to account for the true neural information about task variables. Hence we named it “normalized distance” (ND). This metric is similar to the one used to characterize the community structure in fMRI data (Schapiro et al. (2016)), as well as the “abstraction index” used in analyzing electrophysiological data (Bernardi et al. (2019)). Both of these metrics and ND describe the community structure of neural states organized by task variables. The metrics used in Schapiro et al. (2016) and Bernardi et al. (2019) are computed by dividing the across-group correlation coefficient or Euclidean distance with the same quantity within group. The difference between ND and the previous two metrics is that ND is computed on a moment-by-moment basis. Therefore it reveals the dynamics of the neural information.

When analyzing recordings from a population of neurons, it is often convenient to represent the simultaneous activity of all neurons as a point in a “state space”. A state space is a high dimensional space where each axis represents the activity (in our case binned spike counts) of one neuron. Over time, the population dynamics can be represented by a trajectory through the state space.

The ND for two task conditions *A* and *B* is a function of time ND(*A, B, t*). At each point in time, it is defined to be the average Euclidean distance between pairs of neural trajectories that represent different task conditions divided by the average Euclidean distance between pairs of neural trajectories that represent the same task condition (see Figure 2 for a graphical illustration).

**Figure 2:**
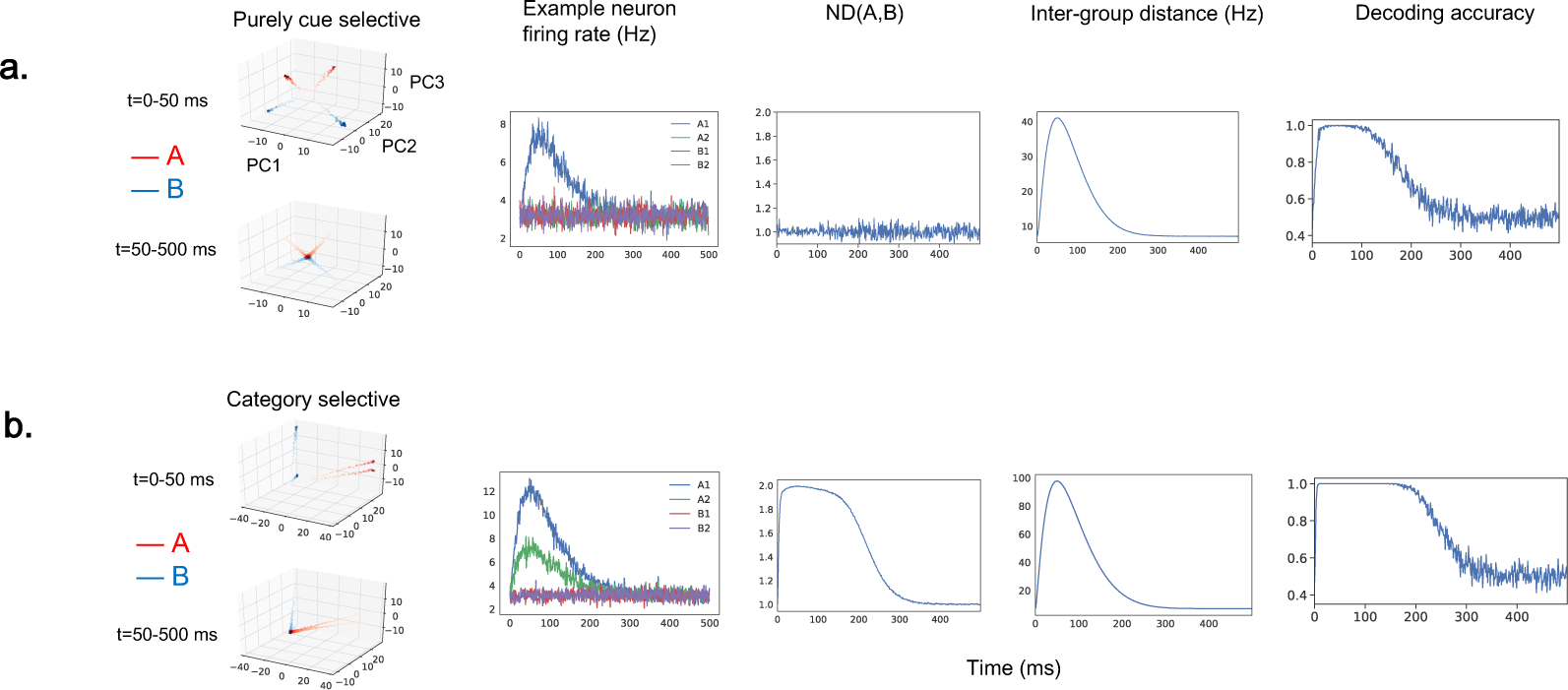
Normalized distance (ND) characterizes the geometry of neural trajectories in state space. We simulated an experiment where a neural population responds to one of the four cues during each trial. We compared different metrics for extracting information about stimulus category in two simulated neuron populations. One population contain neurons that are purely selective for one of the four cues (A1, A2, B1 or B2) (cue-selective population, **a.**). The other population contain neurons with selectivity for both the cue category (A,B) and the cue identity (A1, A2, B1, B2) (category-selective population, **b**). (Left) Low-dimensional representation of the neural trajectories for both populations in the top 3 principle component space. Each trajectory represents the trial-averaged population activity under one condition. (Middle left) The condition-averaged firing rates for an example neuron in the population. The example neuron in the cue-selective population fires most for A1 and remains baseline firing for all the other conditions (top). The example neuron in the category-selective population fires most for A1, less for A2 and remains at baseline firing rate for B1 and B2 (bottom). (Middle) ND for the category-selective population goes above 1, and falls back when the single cell selectivity returns to baseline. On the other hand, ND for the cue-selective population fluctuates around 1. (Middle right) The inter-group distance (the numerator of ND) for both the cue-selective and category-selective populations goes up due to the increased overall firing rate with time in both populations. Therefore the inter-group distance alone, without proper normalization does not accurately reflect the neural information. (Right) The decoding accuracy for cue category (A vs. B) quickly grows above chance in both populations due to larger raw distances between all pairs of neural trajectories. Therefore, although category information is only explicitly encoded in the category-selective population, the decoding accuracy does not reflect this distinction.

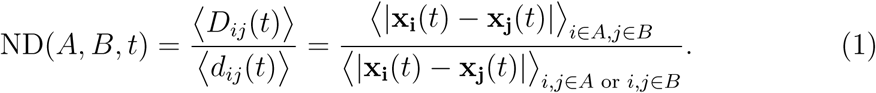

In the equation above, *i* and *j* are indices for neural trajectories, *D*_*ij*_ and *d*_*ij*_ represent the Euclidean distance between neural trajectories *i* and *j* when they belong to the same or different task conditions, respectively. **x_i_**(*t*) and **x_j_**(*t*) are population vectors at time *t* for neural trajectories *i* and *j*. Brackets denote averaging, and vertical lines denote the magnitude of a vector.

ND provides a way of computing the information explicitly encoded in the population from the geometry of the neural data. In Equation 1, the numerator is the average variability in the population code induced by the task condition of interest (here *A* and *B*). The denominator is the average variability induced by all the other factors when the task condition of interest is fixed. The variability in the denominator could come from nested task conditions within *A* and *B*, trial history, or simply intrinsic neuronal noise. An ND(*A, B*) greater than 1 indicates that the population code carries information about task condition *A* versus *B*, because there is extra variability in the population code caused by the difference between *A* and *B* beyond the variability caused by all the other factors. Geometrically, it means that the neural states at time *t* are clustered according to task conditions *A* and *B*. On the other hand, an ND(*A, B*) close to 1 indicates that the population code does not carry information about *A* versus *B*, because the neural states are randomly distributed in the state space.

It is worth emphasizing again that the normalizing part in ND (the denominator in Equation 1) is crucial. A large Euclidean distance between two neural trajectories (the numerator of Equation 1) does not necessarily indicate that the neural population encodes information about the experimental variable. To illustrate this, we simulated two populations of neurons that respond to one of the four cues A1, A2, B1 and B2 (Figure 2) on a given trial. In this case the four cues are organized by two higher-level “categories” A and B. The first population only has selectivity for the identity of the cues (Figure 2a, cue selective population). The second population of neurons has selectivity for both the higher-order categories (whether a cue belongs to category A or B as well as the lower-order cue identity (Figure 2b, category-selective population). The firing rate at each time point was simulated as a Gaussian random variable with a time-varying mean. There are equal number of neurons with a given selectivity in both populations. We computed the normalized distance between the two categories ND(*A, B, t*) as well as the raw distance between the neural trajectories for A and B (inter-group distance, the numerator of ND(*A, B, t*)). As a result, the ND(*A, B, t*) for the two populations have distinctively different time courses that correlate with their single-cell selectivity for the higher-order variables (Figure 2, middle panel). On the other hand, the inter-group distance between A and B is the same for both populations, showing that the raw distance is not enough to capture the information content in the population (Figure 2, middle right panel). The neural trajectories used in the computation of ND could be the population activity during a single trial, or trial-averaged activity for one task condition. This is largely a practical choice and does not affect the rationale above. For this dataset the neural trajectories represent the trial-averaged neural activity over time.

### 2.3 Comparison with decoding accuracy and single cell percentage of explained variance

Another way of quantifying the information in the population code is by constructing a pattern classifier to separate neural activity vectors in the state space according to task conditions. However decoding may miss important geometric properties of the neural states. To show this, we trained a linear discriminant analysis (LDA) classifier to distinguish between cue category A and B using the two simulated neural populations above. As shown in Figure 2 (rightmost panel), the decoders for the two populations behave almost identically. They can both decode the cue category perfectly after a certain point in time. Therefore, decoding accuracy can be blind to important geometric charateristics in the neural data.

The form of the normalized distance also reminds one of single cell measures such as percent of explained variance (PEV). However, computing PEV in the case of nested task conditions can be tricky. In the example above, one cannot compute PEV by simply constructing a linear model for the firing rate of individual neurons using all stimuli and category conditions (A1, A2, B1, B2, A, B) as regressors as the regressors will be correlated. Another way would be to use only the cue category (A, B) as regressors, but in this way the purely cue-selective neurons would also have non-zero PEV for the category. One can construct auxiliary regressors as in Brincat and Miller (2016) to balance out the PEVs for different regressors, which is reminiscent of the normalizing term in the computation of ND. However this technique can not generalize to situations when there are unequal number of cues that belong to one category, whereas ND is generalizable to any situation involving nested task conditions.

## 3 Results

We applied the normalized distance metric introduced above to compute the time evolution of the information encoded in the neural population. We looked at the encoding of every task condition in the task, namely the identity of the cues (*A*_1_ vs. *A*_2_ vs. *B*_1_ vs. *B*_2_), associates (*A*ʹ vs. *B*ʹ), the category of the cues (if the cue belongs to associate A or B) and decision/movement preparation (match vs. non-match).^1^

We found that both PFC and HPC populations encode the identity of the cue and associate stimuli. We also found the neural information in PFC is more persistent compared with HPC. Moreover, there is intermittent information about cue category in PFC but not in HPC. In one of the two monkeys, the PFC population also show a slowly ramping decision/movement information at the end of the trial.

To look at learning-dependent change in neural activity, we divided the learning into 3 stages (early, mid, late) by evenly dividing all trials within a session. Across learning, we found in one of the two monkeys that the neural information about decision/movement preparation in PFC gradually increases. We also found in the same monkey that the neural information about the identity of the associate stimuli increases with learning in HPC.

Neural trajectories were obtained from condition-averaged firing rates. Firing rates were computed every 1 ms using a moving window of 50 ms. Therefore there are 4 neural trajectories during the cue period (4 cues) and 8 neural trajectories after the associate stimulus is presented (4 cues × 2 associates). A given ND was computed by partitioning all neural trajectories into groups that correspond to each task condition, and we then normalized the average across-group distance with average within-group distance, as detailed in Section 2.2. For the all population analysis except for the one on trial outcome (Section 3.5), only sessions where 5 or more neurons were simultaneously recorded were used in computing ND, and for a given session only correct trials were included in calculating the condition-averaged response. For the trial outcome analysis, all sessions are used. All the analysis were performed using simultaneously recorded neurons. To test which part of the trial the ND is significantly larger than 1, we used cluster-based permutation test (Maris and Oostenveld (2007)). To perform this test, we first created a dataset representing ND at chance level which is 1 across all time points. The sample size of the chance dataset is the same as the actual data. As a first-order test, we computed for each time point a one-sided t-test against the chance. The cluster-based permutation test then stipulates a cluster of time points as having an ND significantly larger than chance when the sum of the t-statistic within that cluster is larger than 95% of the random shuffles. In the subsequent sections the time evolution of ND for each task condition across learning stages and brain areas will be presented in turn.

### 3.1 Cue encoding

The ND among the 4 cue stimuli was computed as a function of time. The normalization factor in Equation 1 was calculated from the distances between trajectories encoding the same cue but different associate stimuli (Figure 3a). According to the discussion above, an ND larger than 1 implies that the population contains information about the identity of the cues. On the other hand, an ND close to 1 implies that the population does not carry information about the cues.

**Figure 3:**
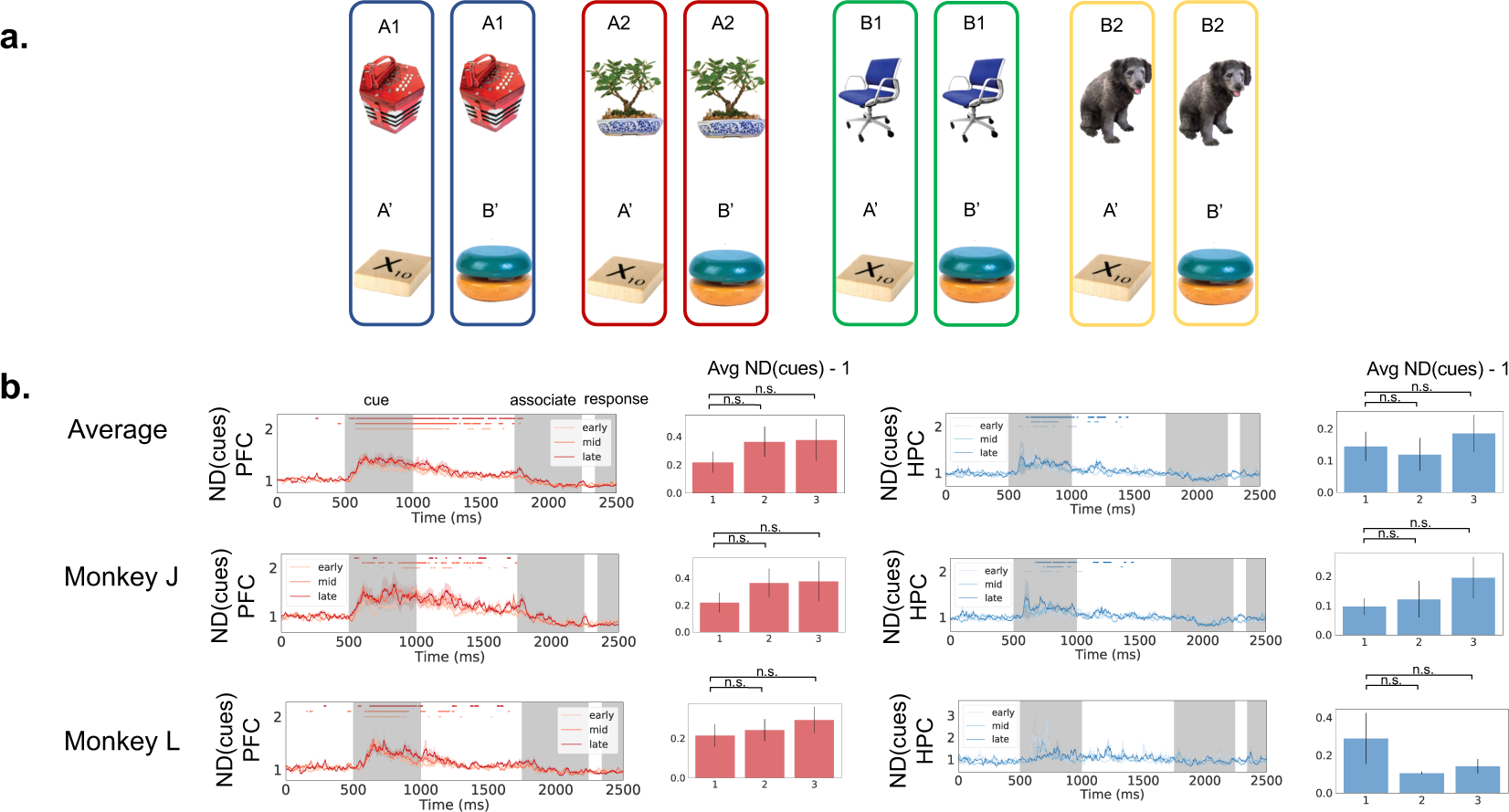
Normalized distance for cues. To calculate the ND among cues. trial-averaged neural trajectories were grouped according to the cue identity (**a**, different colors correspond to different groups). ND(cues) was then the ratio of average Euclidean distances between neural trajectories across groups with that of neural trajectories within groups. Both PFC (**b**) left and HPC (**b** right) encode cue information. The information in PFC persists longer, potentially reflecting the longer timescale of the neuronal activity in PFC compared to HPC. The average ND during the cue presentation does not change with learning in both areas. Shaded area shows 68% confidence interval computed from 10000 iterations of bootstrap resampling across sessions. Dots on top represent timepoints when the ND is significantly larger than 1 (*p <* 0.05, cluster-based permutation test). Gray shaded areas indicate time periods when the cue and associate are presented and response is made.

Figure 3 shows the ND among cues as a function of time. In both PFC and HPC, ND(cues) rapidly goes up when the cue is presented at t=500ms, reflecting the encoding of the cue information at stimulus presentation. The information about the cues then gradually decays to baseline. For PFC, it rapidly decreases after the subsequent stimulus (associate) is presented. Notably, PFC has a much more sustained ND than HPC. This implies that the network timescale in PFC may be longer than HPC. There are no significant learning-dependent changes in ND for both PFC and HPC during the cue period (Fig 3b, Mann-Whitney U test). Overall, the results show that both PFC and HPC populations carry transient information about the cues, and that information lasts longer in PFC.

### 3.2 Associate encoding

The ND between the 2 associate stimuli ND(A’,B’) was computed as in Equation 1, where the denominator is the average distance between neural trajectories encoding the same associate stimuli but different cue stimuli (Figure 4a). The ND was only computed after the associate was presented. As shown in Figure 4, the ND between the two associates (A’ vs. B’) rapidly increases when the associate is presented (t=1750ms). Similar to cue encoding, the ND in PFC shows sustained information for a longer period of time than that in HPC (compare Figure 4b, red and blue lines). In addition, we observed an interesting learning-dependent change in one of the monkeys: the average ND during the associate presentation period is significantly smaller during early learning period (the first 1/3 of the trial) than mid (the middle 1/3) and late (the last 1/3) learning periods (Figure 4d, Mann-Whitney U test, *p* = 0.003 between early and late. *p* = 0.01 between early and mid). This effect persists when the analysis was done on the combined data from both monkeys. This possibly reflects some reconfigurations within the HPC circuit that enable it to represent the associate stimulus more strongly with learning.

**Figure 4:**
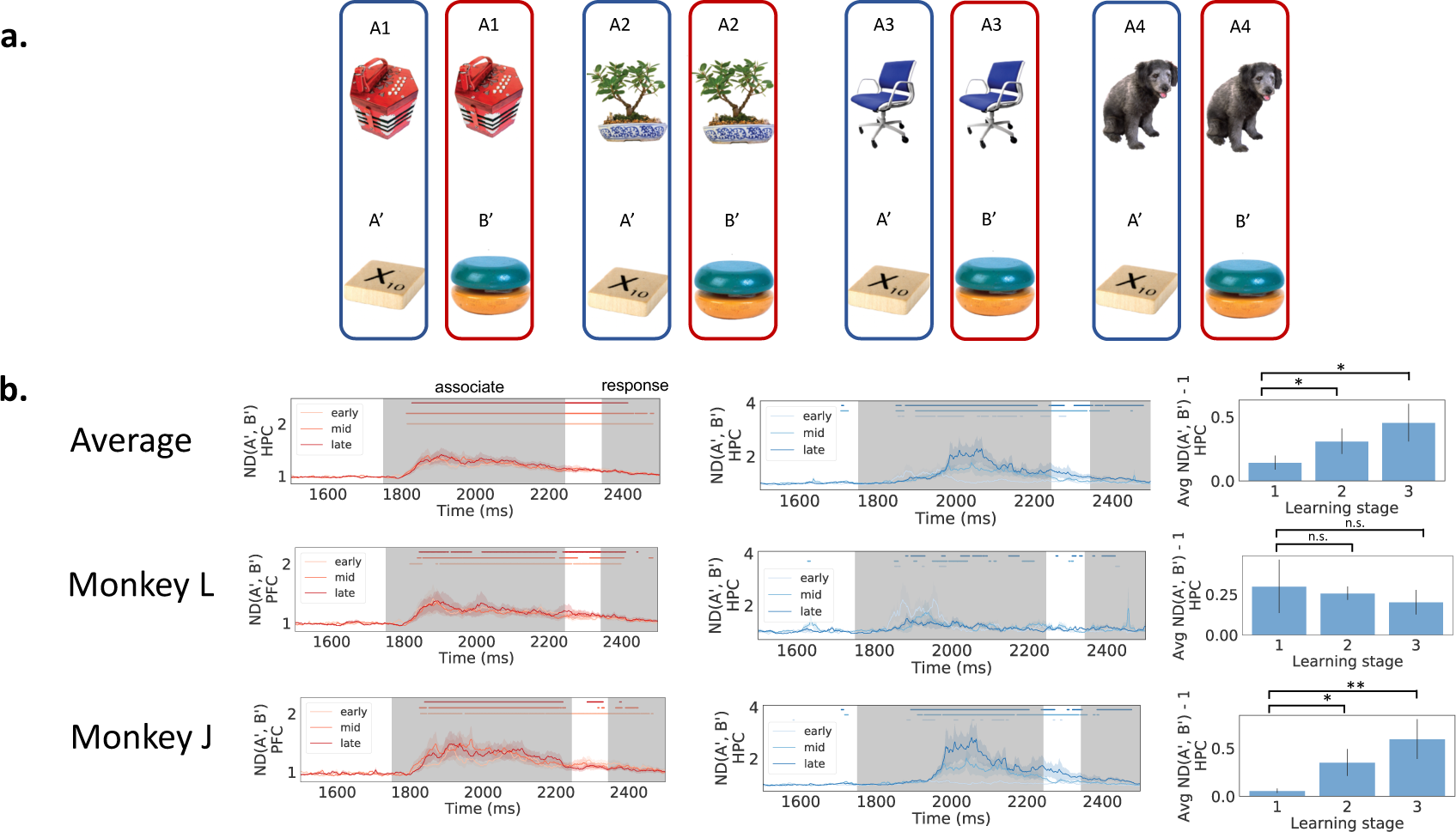
Normalized distance for associates. The ND for associates was computed during the period after the associate has been presented. Neural trajectories were grouped according to the identity of the associate stimuli (**a**). Both PFC (**b** left) and HPC (**b** right) encode information about the identity of the associate stimulus. The information in PFC sustains longer than HPC (Shaded area shows 68% confidence interval computed from 10000 iterations of bootstrap resampling across sessions. Dots on top represent timepoints when the ND is significantly larger than 1 (*p <* 0.05, cluster-based permutation test). Gray shaded areas indicate time period when the cue and associate are presented and response is made). In the HPC of monkey J, the information about the associate stimulus is stronger later in learning (rightmost panel of **b**, ND averaged across the associate presentation period. Asterisks indicate statistical significance using Mann-Whitney U test).

### 3.3 Category information encoding

We next investigated whether the information about the cue category was encoded in PFC and HPC, by which we mean whether the cue was paired with associate A’ or B’. We define cue A1 and A2 to be in cue category A, and B1 and B2 to be in cue category B. The ND between cue categories (ND(A,B)) was computed according to Equation 1. Different neural trajectories were grouped according to the category of their cues (Figure 5a). Importantly, since there are 4 task conditions before the associate was presented and 8 after, the grouping of neural trajectories were different before and after the associate was presented as well. Therefore, ND(A,B) was calculated in different ways for the two stages (Figure 5a).

**Figure 5:**
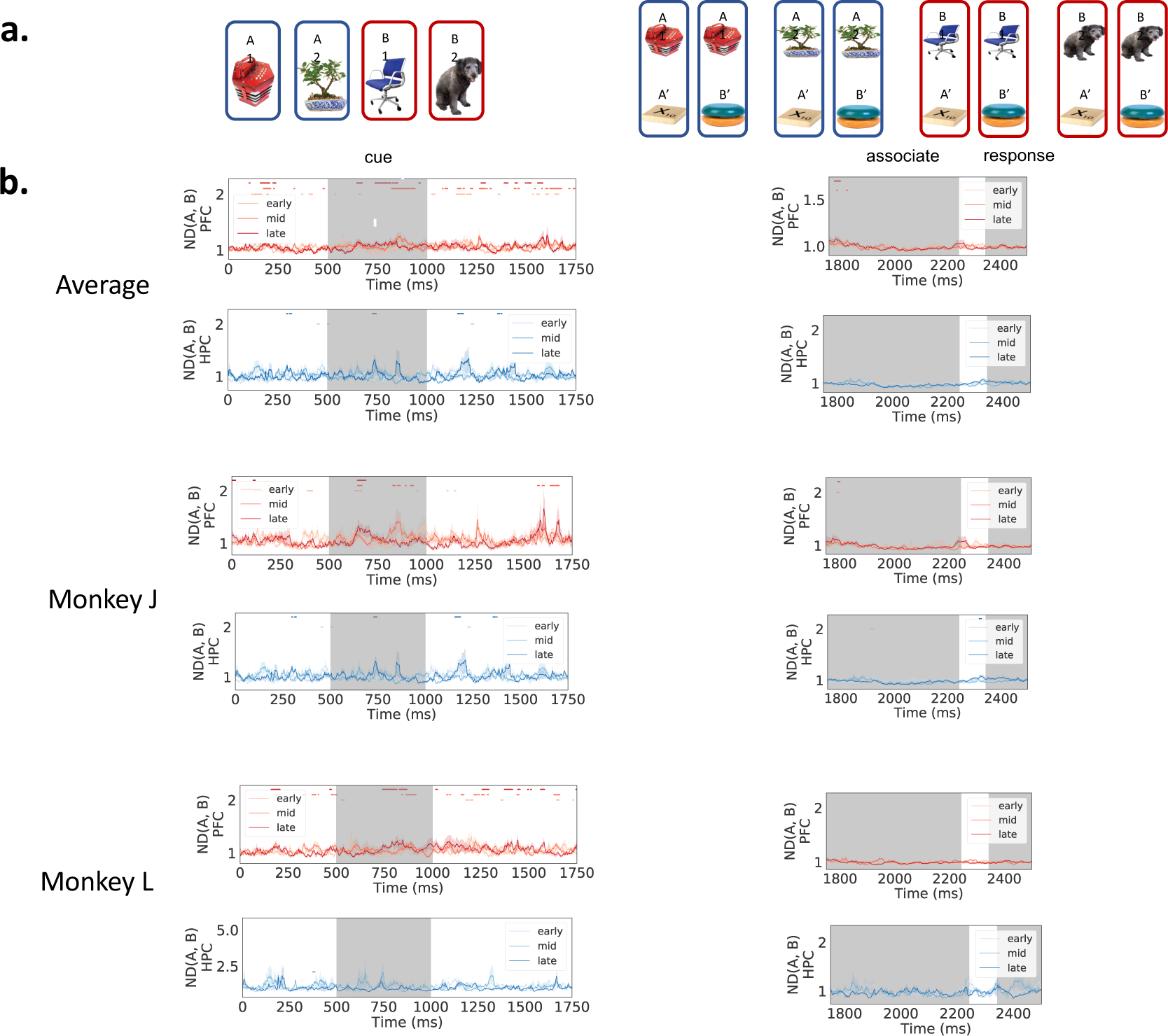
Normalized distance for categories. The ND for cue category was calculated by grouping the neural trajectories according to the cue category (**a**). PFC has intermittent information about the category information (that *A*_1_ and *A*_2_ both predict *A*ʹ, and *B*_1_ and *B*_2_ predict *B*ʹ, **b**, red lines), whereas HPC did not encode this information (**b**, blue lines). Shaded area shows 68% confidence interval computed from 10000 iterations of bootstrap resampling across sessions. Dots on top represent timepoints when the ND is significantly larger than 1 (*p* < 0.05, cluster-based permutation test). Gray shaded areas indicate time periods when the cue and associate are presented and response is made.

As shown in Figure 5, in PFC, there is intermittent information about the cue category during the cue and delay periods (Figure 5b, left). This might reflect some representation for the cue category. However, this signal is much weaker than the information about cues (compare Figure 3b), indicating that neural trajectories are mainly organized by cues and only weakly by the cue category. On the contrary, ND(A,B) for HPC fluctuates around 1 throughout the trial, indicating that HPC does not encode information about the category of the cue.

### 3.4 Decision variable/movement encoding

To investigate the neural signals about the upcoming decision/movement after the monkeys received the sequence of stimuli, we computed the ND between match and non-match trials. Since we only looked at correct trials, decision about match versus non-match is perfectly confounded with movement preparation. In this experiment there is no way to look at one of them without the influence of the other.

The ND between match and non-match trials is shown in Figure 6. It was calculated by grouping the neural trajectories according to whether it is a match or non-match trial (Figure 6a). In the PFC of one of the monkeys, the ND between match and non-match trials starts to ramp up halfway during the second stimulus presentation period when all the information needed to form the decision is present. This effect is also present when combining data from both monkeys. There is a latency of about 150 ms between the appearance of the associate information in PFC and that of the decision (compare Figure 6b with Figure 4b). This potentially indicates the time that the monkey took to compare the incoming associate stimulus with the cue stimulus in the working memory. In contrast, in HPC the ND between match and non-match trials fluctuates around 1 until the time of the response saccade (Figure 6f). Therefore HPC does not encode the decision variable during the time when the match/non-match decision is presumably being made.

**Figure 6:**
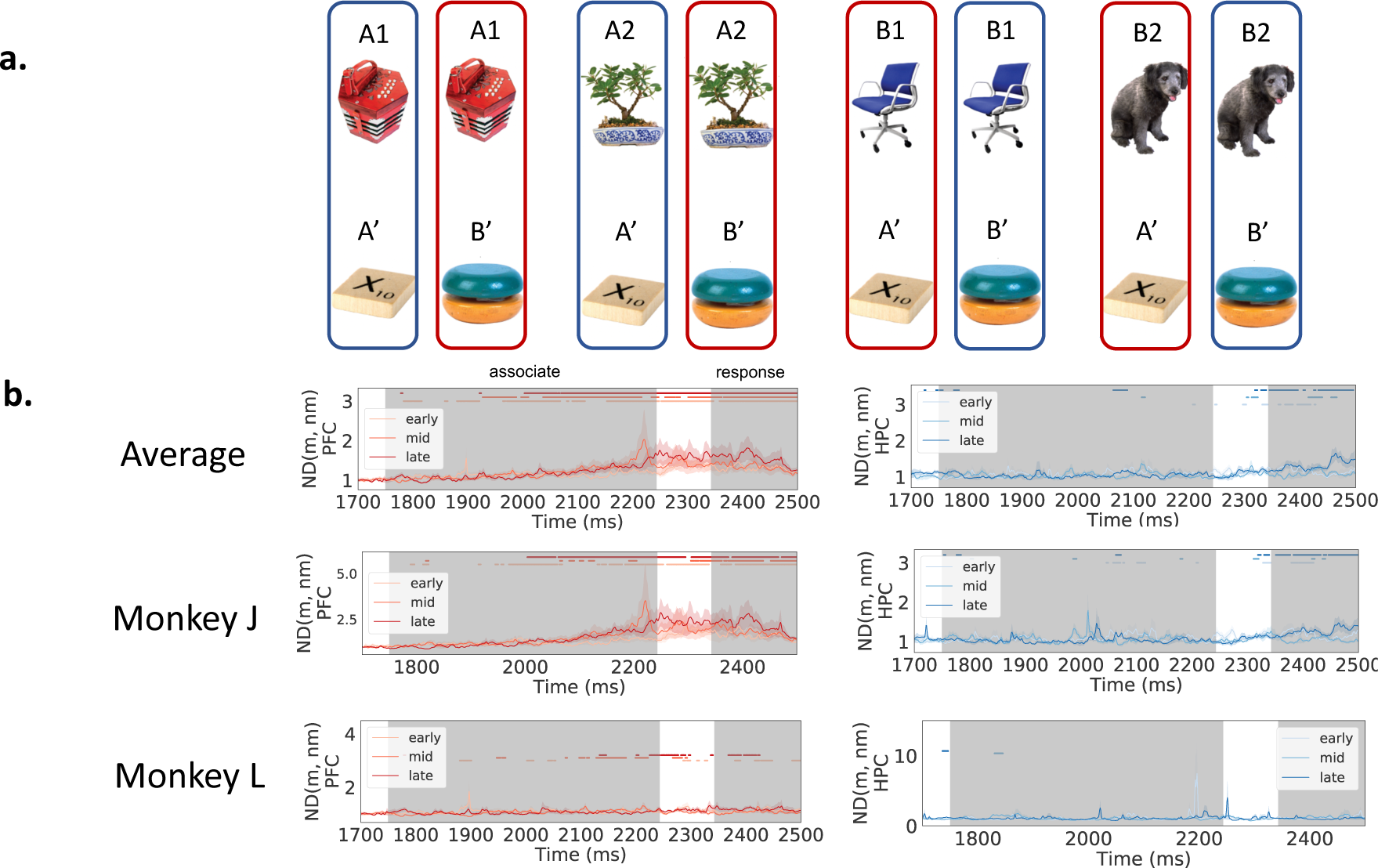
Normalized distance for decision/movement. The ND for decision/movement was calculated by grouping the neural trajectories according to whether it is a match or non-match trial (**a**). In the PFC of monkey J, information about the match versus non-match trial type starts to ramp up halfway through the second stimulus interval (**b**, left). The ND for HPC fluctuates around 1 for both monkeys (**b**, right). Shaded area shows 68% confidence interval computed from 10000 iterations of bootstrap resampling across sessions. Dots on top represent timepoints when the ND is significantly larger than 1 (*p <* 0.05, cluster-based permutation test). Gray shaded areas indicate time periods when the associate is presented and response is made.

### 3.5 Trial outcome encoding

In Brincat and Miller (2015), it was shown that the synchrony between PFC and HPC increases after the trial outcome, potentially serving as the communication signal between the two brain regions to facilitate associative learning. In this section we compute the ND between the rewarded and nonrewarded trials after the trial outcome (Figure 7a). The information of trial outcome persists for seconds in both PFC and HPC, as measured by an ND that is significantly greater than 1. In PFC the ND is significantly greater than 1 throughout the 2.8s interval we looked at after the trial outcome (Figure 7b, red lines). In HPC the ND falls back to 1 more quickly, but the information on trial outcome still persists for more than 1 second (Figure 7b, blue lines). These results show that aside from the synchronous LFPs between PFC and HPC (Brincat and Miller (2015)), the spiking activity in both regions also carries information about the outcome of the previous trial for seconds long ^2^.

**Figure 7:**
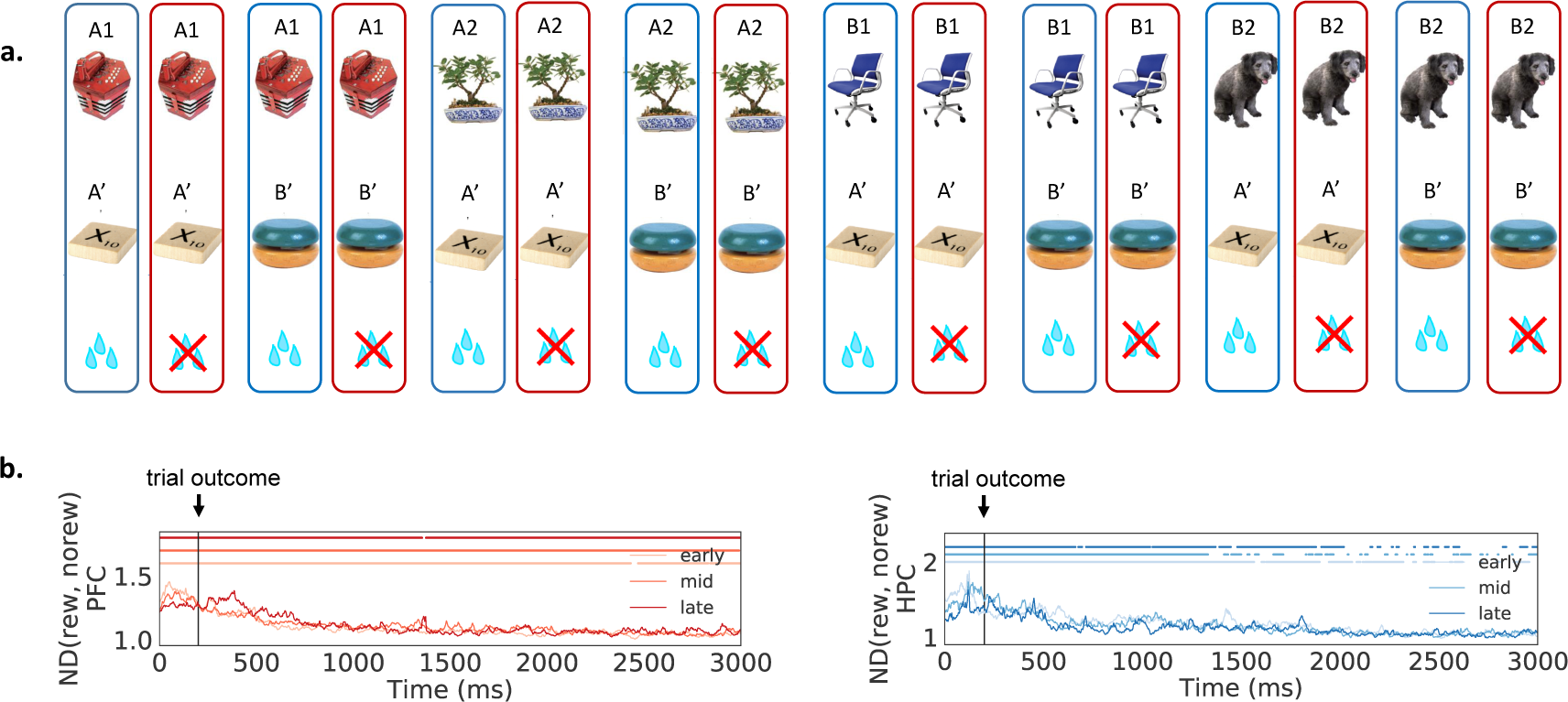
Normalized distance for trial outcome. The ND for trial outcome was calculated by grouping the neural trajectories according to if the animal was rewarded **a**. In PFC (**b**, red lines) and, the information for trial outcome persists throughout the 2.8 seconds, as reflected by an ND significantly greater than 1 (dots on the top part of the figure). In HPC the information about trial outcome also persists for over a second (**b**, blue lines). Shaded area shows 68% confidence interval computed from 10000 iterations of bootstrap resampling across sessions. Dots on top represent timepoints when the ND is significantly larger than 1 (*p <* 0.05, cluster-based permutation test).

## 4 Discussion

In this paper we developed a new metric called normalized distance (ND) that characterizes neural information in the population code based on the geometric organization of neural trajectories in the state space. We then applied this metric to recording from a paired-associate task (Brincat and Miller (2015, 2016)) to compare the dynamics of the coding of different task variables across learning stages and brain areas (PFC and HPC). We found that 1) PFC and HPC both code for the identity of both the cue and the associate stumuli. 2) PFC exhibits a longer network timescale for stimulus coding (Figure 3, 4). 3) The information about the cue category is intermittent in PFC but not significant in HPC (Figure 5). 4) The information about the previous trial’s outcome persists for seconds in both PFC and HPC (Figure 7), We also found evidence in one of the two monkeys that 5) Coding for the associate object increases with learning in HPC but not PFC (Figure 4), and 6) Information for decision/movement slowly ramps up in PFC but not HPC (Figure 6). These effects persist when averaged across both animals.

The results 2, 4, 5, 6 above were not reported in the original paper (Brincat and Miller (2015, 2016)). In particular, the network timescale for object coding was found to be longer in PFC than HPC, potentially providing suitable temporal dynamics for PFC to integrate incoming information from other cortices. It was indeed reported that single neurons in PFC have, on average, longer time constants than motor areas (Murray et al. (2014)).

In the original paper (Brincat and Miller (2015, 2016)), the authors used another metric, bias-corrected percentage of explained variance, to calculate neural information as a function of learning. They discovered that the object category coding was present in PFC but not HPC (Brincat and Miller (2015, 2016)). We find some evidence that corroborate this finding (Figure 5). However, they also found that the object category coding increased with learning, which we did not find using the ND metric. It could be that the different metrics used caused different results, but it should be noted that learning effects are subtle in this experiment.

There are other studies that reported single cell activity during associative learning (Sakai and Miyashita (1991); Suzuki (2007)). These earlier studies used metrics based on single cell firing rates to correlate with learning performance and experimental conditions. In contrast, in this paper the ND metric we used characterize the distributed information in the population code.

The ND metric developed in this paper serves a similar role as decoding accuracy commonly adopted in analyzing population level data. In decoding analysis hyperplanes are constructed that separate training data from different categories as well as possible according to some objective function, and the decoding accuracy reflects how well these hyperplanes separate the held-out test data. It is noted however that in high dimensional cases where the number of neurons is comparable with the number of data points (as in our case here as well as data obtained by modern large-scale recording techniques), the decoding accuracy can be generically high (Buonomano and Maass (2009)) and thus does not necessarily reflect the underlying geometry of the neural code. It is therefore difficult to interpret results obtained by directly comparing decoding accuracies across different recording sessions. On the other hand, ND is directly computed from the geometrical configuration of the data, and therefore can be used regardless of the number of data points compared with the number of dimensions and provide a clear geometrical picture of the underlying neural code, as illustrated using the simulated data in Section 2.3.

In this data set HPC does not encode the category information of the cues that would help predict the upcoming stimuli even after learning. This seems contradictory with the study by Stachenfeld et al. showing that HPC contains a predictive map of the environment (Stachenfeld et al. (2017)) and the study by McKenzie et al. showing rodent HPC population encodes the hierarchical structure of the task (McKenzie et al. (2014)). However, the scenario studied by Stachenfeld et al. is a reinforcement learning task in a spatial setting where sensory experience is almost continuous. The task studied by McKenzie et al. also has a spatial component, and the population activity was observed to be largely organized by spatial context. On the other hand, the task we analyzed here is a simple sequential associative learning task. It could be that in this simple setting, the HPC is not utilized to form more sophisticated relational representations (Eichenbaum (2017))

The normalized distance is directly calculated from the geometry of neuronal population responses in the state space. Therefore it provides a characterization of the degree of “tangling” of the underlying neuronal manifolds. Disentangled neuronal manifolds were argued to be crucial in forming a “good” neuronal representations for higher level processing (DiCarlo and Cox (2007)). An ND larger than 1 indicates that the neuronal manifolds representing different variables are some-what disentangled. However, we do note that other geometric quantities such as curvature (Bernardi et al. (2019); Chaudhuri et al. (2019)) are needed to provide a complete characterization of the neuronal manifold.

## Acknowledgements

This work was supported by ONR MURI award N00014-16-1-2832 (Y.L., M.E.H., E.K.M.), ONR DURIP award N00014-17-1-2304 (Y.L., M.E.H., E.K.M.) and NIMH R37MH087027 (S.L.B., E.K.M.).

## Footnotes

1 In this experiment we cannot tease apart the neural information for decision variable versus general preparation of an upcoming movement.

2 In this analysis we did not look at each monkey individually because there is not enough usable data for one of the monkeys

